# *TMEM106B* and *CPOX* are genetic determinants of cerebrospinal fluid Alzheimer’s disease biomarker levels

**DOI:** 10.1101/2020.05.31.125633

**Authors:** Shengjun Hong, Valerija Dobricic, Isabelle Bos, Stephanie J. B. Vos, Dmitry Prokopenko, Betty M. Tijms, Ulf Andreasson, Kaj Blennow, Rik Vandenberghe, Silvy Gabel, Philip Scheltens, Charlotte E. Teunissen, Sebastiaan Engelborghs, Giovanni Frisoni, Olivier Blin, Jill C. Richardson, Regis Bordet, Alberto Lleó, Daniel Alcolea, Julius Popp, Christopher Clark, Gwendoline Peyratout, Pablo Martinez-Lage, Mikel Tainta, Richard J. B. Dobson, Cristina Legido-Quigley, Kristel Sleegers, Christine Van Broeckhoven, Rudolph E Tanzi, Mara ten Kate, Michael Wittig, Andre Franke, Frederik Barkhof, Simon Lovestone, Johannes Streffer, Henrik Zetterberg, Pieter Jelle Visser, Lars Bertram

## Abstract

**Background:** Neurofilament light (NF-L), chitinase-3-like protein 1 (YKL-40), and neurogranin (Ng) are utilized as biomarkers for Alzheimer’s disease (AD), to monitor axonal damage, astroglial activation, and synaptic degeneration, respectively. Here we performed genome-wide association study (GWAS) analyses using all three biomarkers as outcome.

**Methods:** DNA and cerebrospinal fluid (CSF) samples originated from the European Medical Information Framework AD Multimodal Biomarker Discovery (EMIF-AD MBD) study. Overlapping genotype/phenotype data were available for n=671 (NF-L), 677 (YKL-40), and 672 (Ng) individuals. GWAS analyses applied linear regression models adjusting for relevant covariates.

**Findings:** We identify novel genome-wide significant associations with markers in *TMEM106B* and CSF levels of NF-L. Additional novel signals were observed with DNA variants in *CPOX* and CSF levels of YKL-40. Lastly, we confirmed previous work suggesting that YKL-40 levels are regulated by *cis* protein quantitative trait loci (pQTL) in *CHI3L1*.

**Interpretation:** Our study provides important new insights into the genetic architecture underlying inter-individual variation in all three tested AD-related CSF biomarkers. In particular, our data shed light on the sequence of events regarding the initiation and progression of neuropathological processes relevant in AD.

## Background

Elucidation of the genetic architecture underlying Alzheimer’s disease (AD) susceptibility has recently seen substantial progress owing to the application of large-scale analysis approaches in the context of genome-wide association studies (GWAS). Based on results from the two most recent and largest GWAS in the field ^1,2^, there are now more than 30 independent loci showing genome-wide significant association with AD risk^3^. In contrast, the genetic underpinnings determining inter-individual variation in levels of molecular AD biomarkers are less well known. Apart from the two “core” proteins, i.e. amyloid β 42 (Aβ42) and tau, the aggregation of which represents the neuropathological plaque and tangle hallmarks of the disease, there are currently only very few GWAS shedding light on the genetic factors determining blood or cerebrospinal fluid (CSF) biomarkers levels in AD. In an effort to close this knowledge gap, we combined CSF and genome-wide single-nucleotide polymorphism (SNP) genotyping data generated in the European Medical Information Framework AD Multimodal Biomarker Discovery study (EMIF-AD MBD) ^4^ and performed the first *bona-fide* GWAS on CSF levels of neurofilament light chain (NF-L), chitinase-3-like protein 1 (YKL-40), neurogranin (Ng), reflecting axonal damage, astroglial activation, and synaptic degeneration, respectively.

NF-L is one type of four different neurofilament subunits which function as structural components of the neural cytoskeleton ^5^ performing essential roles in axon development ^6^ and synaptic function ^6^. As such, NF-L is considered one of several “core” biomarkers of axonal injury and neurodegeneration across neurological diseases ^7,8^. In addition, other recent data suggest that changes in NF-L serum levels can predict disease onset and progression of brain neurodegeneration at very early, pre-symptomatic stages of familial AD ^9^. YKL-40 is a glycoprotein produced in several inflammatory conditions and cancers ^10^, and was classified as an “emerging” AD biomarker in a recent meta-analysis ^7^. While its precise physiological role remains elusive, it appears that in AD, YKL-40 is predominantly expressed in astrocytes and likely plays a role in the inflammatory response occurring near Aβ plaques ^10,11^. Finally, Ng is a neuron-specific protein, mainly expressed in the cortex and hippocampus, where it is involved in synaptic long-term potentiation and learning ^12–14^. In AD, CSF Ng was proposed to represent a marker of synaptic degeneration and was recently reported to correlate with cognitive decline ^15^.

The results of our GWAS identified novel genome-wide significant associations with markers in the established frontotemporal lobe dementia (FTLD) risk gene *TMEM106B* and CSF levels of NF-L in the EMIF-AD MBD dataset. Additional novel signals were observed between DNA variants in *CPOX* and CSF levels of both YKL-40 and Ng. Finally, we detected very strong and highly significant association between markers in *CHI3L1* and CSF levels of YKL-40, representing the only *cis* protein quantitative trait locus (pQTL) in our analyses.

## Methods

### Sample description

The ascertainment procedures for the EMIF-AD MBD dataset are described elsewhere ^16^. In brief, the dataset includes 1221 elderly individuals (years of age: mean = 67.9, SD = 8.3) with different cognitive diagnoses at baseline (NC = normal cognition; MCI = mild cognitive impairment; AD = AD-type dementia) and the clinical follow-up data were available for 759 individuals. Depending on the availability of the clinical records, each phenotype has slightly different effective sample sizes. The demographic information for the three quantitative CSF phenotypes of the EMIF-AD MBD dataset utilized in this study is summarized in Table 1.

**Table 1.** Demographic information and summary of CSF traits used in GWAS analyses.

| **CSF biomarker** | **CSF biomarker description** | **Detection method** | **N** | **N male / female** | **Mean age ± SD (range)** | **N Controls** | **N MCI** | **N AD** |
| --- | --- | --- | --- | --- | --- | --- | --- | --- |
| NF-L | Neurofilament light | ELISA (UmanDiagnostics) | 671 | 319/352 | 69.49±8.35(45.94-92.29) | 122 | 395 | 154 |
| YKL-40 | Chitinase-3-like protein 1 | ELISA (R&D systems) | 677 | 323/354 | 69.42±8.31(45.94-92.29) | 122 | 401 | 154 |
| Ng | Neurogranin | Immunoassay (in-house) | 672 | 319/353 | 69.50±8.34(45.94-92.29) | 122 | 398 | 152 |

### DNA extraction, Genotype imputation and quality control

All participants had provided written consent prior to participation and institutional review board (IRB) approvals for the utilization of the DNA samples in the context of EMIF-AD MBD were obtained by the sample collection sites. A detailed account of the genotyping procedures and subsequent bioinformatic workflows can be found in Hong et al. ^17^. In brief, a total of 936 DNA samples were sent for genome-wide SNP genotyping using the Infinium Global Screening Array (GSA) with Shared Custom Content (Illumina Inc.). After extensive QC and imputation, a total of 7,778,465 autosomal SNPs with minor allele frequency (MAF) ≥0.01 were retained in 898 individuals of European ancestry for the downstream genome-wide association analyses.

### CSF biomarkers

Details of the CSF biomarker measurements can be found in Bos et al. ^4^. In brief, the CSF specimen were collected individually at each of the 11 EMIF-AD MDB participating sites. CSF samples were shipped to Department of Psychiatry and Neurochemistry at University of Gothenburg, Sweden.

Relevant to the analyses presented here, NF-L levels were measured using a commercial ELISA (NF-light ELISA, UmanDiagnostics, Umeå, Sweden ^18^). Ng levels were measured using an in-house immunoassay for Ng ^19^. YKL-40 levels were measured using a human chitinase-3 quantikine ELISA kit (R&D systems, Inc, Minneapolis, MN ^20^). To reduce the skewness of phenotype distributions, data for all three CSF variables were log-transformed prior to analysis (Supplementary Figure 1).

### GWAS and post-GWAS analyses

Linear regression (using mach2qtl ^21,22^) was utilized to perform SNP-based association analyses using imputation-derived allele dosages as independent variables and the log-transformed concentrations of CSF NF-L, CSF Ng, and CSF YKL-40 as dependent outcome variables. Covariates included into the regression models were sex, age at examination, diagnostic groups (coded as AD = 3, MCI = 2, controls = 1), and ancestry-specific principal components (PCs; here the first 5 were used). The genomic inflation factor was calculated in R using the “GenABEL” package^23^. FUMA (http://fuma.ctglab.nl/) ^24^ was used for post-GWAS analyses, including gene annotations and functional mappings. This also included performing gene-based GWAS analyses using MAGMA ^25^ as implemented in FUMA. Genome-wide significance was defined as 5E-08 for the SNP-based and 2.651E-6 for the gene-based analyses (as recommended by FUMA).

### Polygenic risk score (PRS) analysis

Summary statistics of the two largest and most recent AD case-control GWAS ^1,2^ were utilized for calculating PRS for each individual in EMIF-AD MDB. PRS were constructed for 11 different P-value thresholds (i.e 5E-08, 5E-06, 1E-04, 0.01, 0.05, 0.10, 0.20, 0.30, 0.40, 0.50, 1.00) using PLINK v1.9 ^26^ after removal of ambiguous SNPs (A/T and C/G), filtering by imputation quality (minimac3 r^2^ ≤0.8), allele frequency (MAF ≤0.01) and linkage disequilibrium (LD) pruning. The resulting PRSs were utilized as independent variables in the regression models adjusting for sex, age, diagnosis, and PC1-5 as covariates as in the primary GWAS analyses. Variance explained (R^2^) was derived from comparing results from the full model (including PRS and covariates) vs. the null model (linear model with covariates only).

*A detailed description of all methods and procedure applied in this study can be found in the Supplementary Material*.

## Results

### GWAS analyses using CSF neurofilament light (NF-L) levels

The GWAS analyses using CSF NF-L as outcome yielded five SNPs showing genome-wide significant association (Figure 1A, Table 2, and Supplementary Table 1). These SNPs are located in two distinct loci on chromosome 7 (i.e., on chr. 7q36.1 [rs111748411, rs3094407] and 7p15.3 on [rs77589784]), while the other two are located on chromosomes 1p36.12 (rs4654961) and 10q26.3 (rs138898705; Figure 1A, Supplementary Table 1 and Supplementary Figure 2a-b). MAFs for SNPs in all but the 7q36.1 locus were around 1% complicating any inferences and functional interpretations of these variants given the limited size of our dataset. For the two common SNPs in the chr. 7q36.1 locus (i.e., rs111748411, rs3094407), post-GWAS variant annotation in FUMA suggested no obvious functional consequences (Supplementary Table 2).

**Figure 1.**
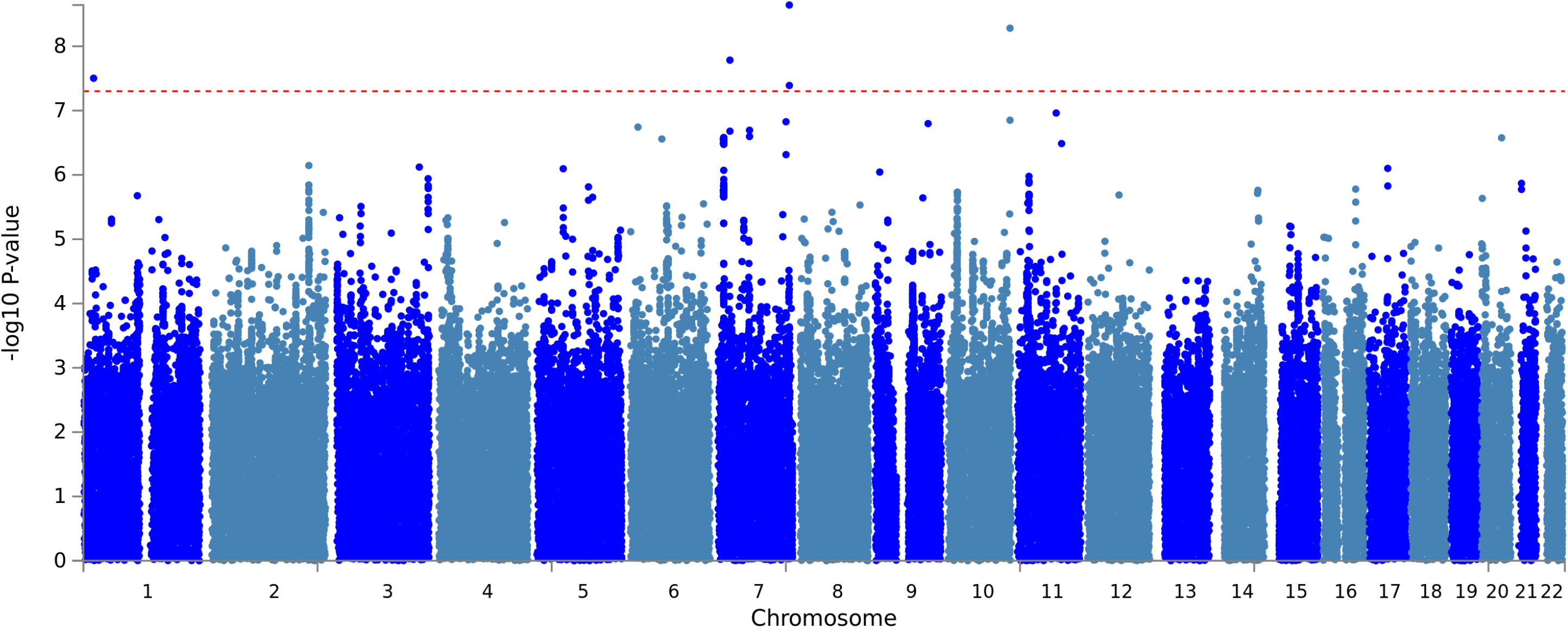
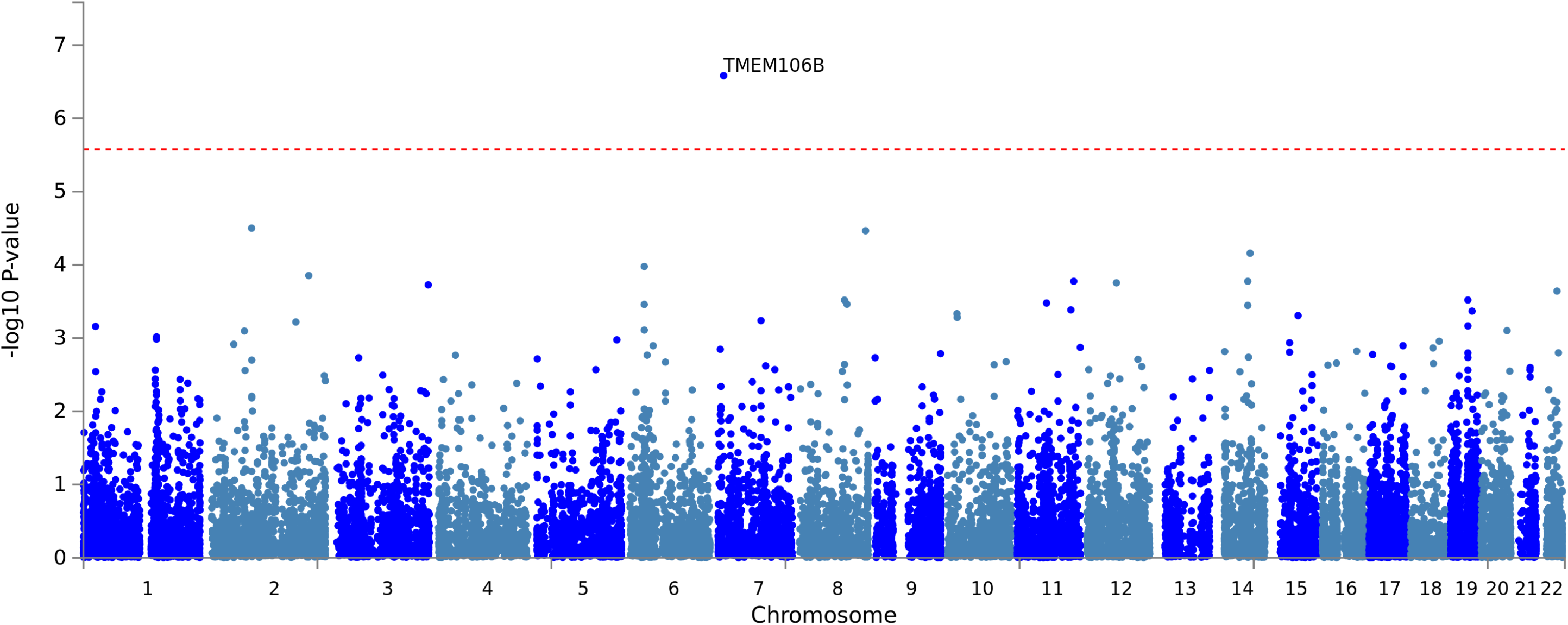
Manhattan plots of **1A)** SNP level and **1B)** gene-level genome-wide association results using log-transformed CSF NF-L levels as outcome trait (n = 671). Gene assignments are according to FUMA ^24^. Dotted red lines represent the threshold for genome-wide significance, i.e. α = 5.0E-08 for SNP-based (1A) and α = 2.651E-6 for gene-based (1B) analyses (see Methods).

**Table 2.**
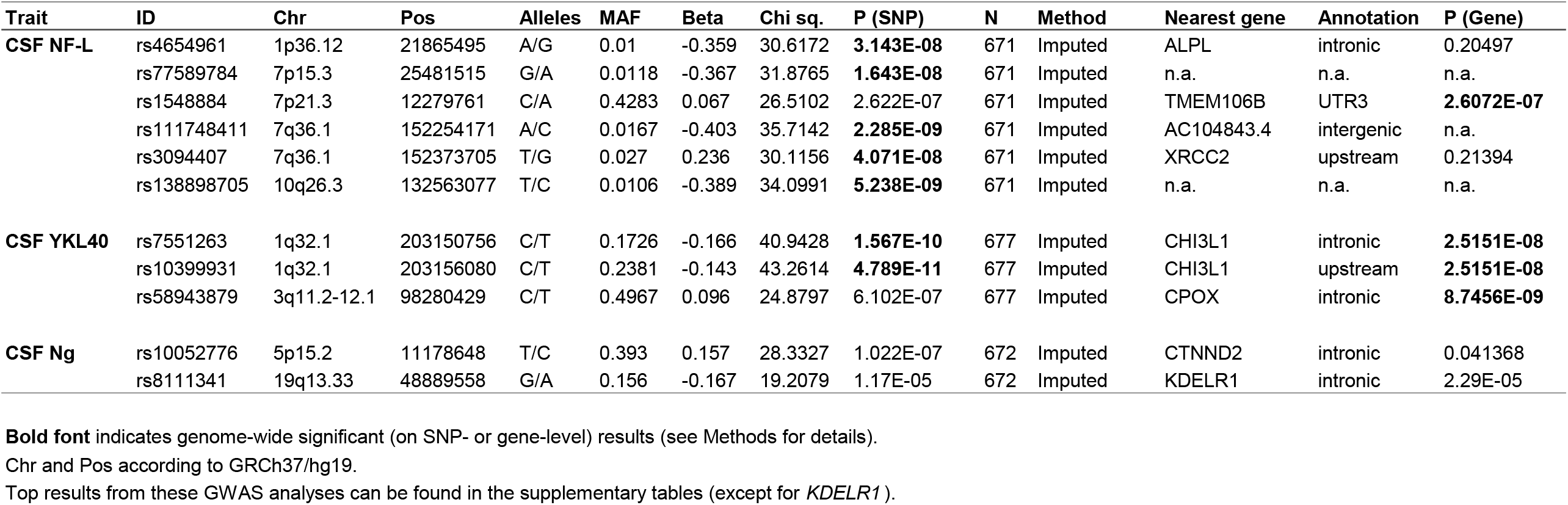
Top results from GWAS analyses of three CSF traits measured in the EMIF-AD MBD dataset.

In contrast to the SNP-based results, the gene-based analyses using MAGMA elicited a third locus on chromosome 7 (7p21.3) at transmembrane protein 106B (*TMEM106B*) showing genome-wide significant association with CSF NF-L levels (Figure 1B, Table 2, and Supplementary Table 1). This gene, which is an established genetic risk modifier for frontotemporal lobar degeneration (FTLD) ^27,28^, contains 187 SNPs of which the majority (n = 124 SNPs) are in strong LD (r^2^ >0.6) with the lead variant in *TMEM106B*, i.e., rs1548884 (single SNP P = 2.62E-07, Table 2 and Supplementary Table 1). FUMA-based functional annotations show one non-synonymous variant (rs3173615; in nearly perfect LD [r^2^>0.99] with rs1548884, Supplementary Tables 2 & 3) eliciting a Thr185Ser change with a CADD score of 21.4, and a predicted “moderate” impact by ENSEMBL’s variant effect predictor (VEP) algorithm. In addition to possibly exerting an effect on protein function by directly altering the aminoacid sequence of *TMEM106B*, the same variant is also reported as a modest expression quantitative trait locus (eQTL) in cortical brain samples of the Genotype Tissue Expression (GTEx, v8) project (P=3.8E-05; Supplementary Table 3). Interestingly, the lead variant in our CSF NF-L GWAS (rs1548884) is in strong LD (r^2^=0.98; Supplementary Table 2) with the SNP originally identified and subsequently replicated to be associated with FTLD (rs1990622, P=3.12E-07, Supplementary Table 1; Van Deerlin VM et al. ^27^).

### GWAS analyses using CSF YKL-40 levels

The SNP-based GWAS using CSF YKL-40 levels yielded one genome-wide significantly associated locus on chromosome 1q32.1. This signal was driven by 3 independent SNPs (i.e., rs7551263, rs1417152, and rs10399931; Figure 2A, Table 2, Supplementary Table 4) and also represents the single most significant GWAS signal of this study (P=4.79E-11 for rs10399931). Unlike the GWAS results for the other two CSF markers analyzed here, the strongest results were observed with relatively common variants showing allele frequencies between ~16 and 21% in people of European ancestry (Supplementary Table 5a). While FUMA-based gene annotations (Supplementary Table 5b) highlight up to 26 different gene symbols in the implicated region, the most obvious candidate of likely biological relevance is *CHI3L1* (chitinase 3 like 1), i.e., the gene encoding YKL-40 protein. In other words, this GWAS results represent a bona fide *cis* protein QTL (pQTL) result, a finding that is also in good agreement with the non-AD literature (see discussion). Furthermore, and corresponding to these pQTL results, eQTL annotations summarized by FUMA converge on *CHI3L1* as the most strongly and most significantly associated gene when using the YKL-40-associated SNPs or their proxies as input. Manual lookup of the top eQTL SNP (rs10399931, i.e. the same as in our GWAS) on the GTEx portal (v8) revealed that this variant functions as a *CHI3L1* eQTL in 24 GTEx tissues, including one brain area (i.e., “Brain - Frontal Cortex (BA9)”, eQTL P = 1.7E-06; Supplementary Table 6). Interestingly, this SNP is also listed as methylation QTL (mQTL) on the mQTL database (http://www.mqtldb.org/) ^29^, a finding corroborated in genome-wide DNA methylation data generated in parallel in the same EMIF-AD MDB dataset (R. Smith, S. Hong, L. Bertram, K. Lunnon, et al., manuscript in preparation). However, since the top associated variant (rs10399931) scores relatively low using *in silico* predictions (e.g., CADD = 0.724, RDB = 1f; Supplementary Table 5), it may not represent the functional entity underlying the observed GWAS and QTL signals.

**Figure 2.**
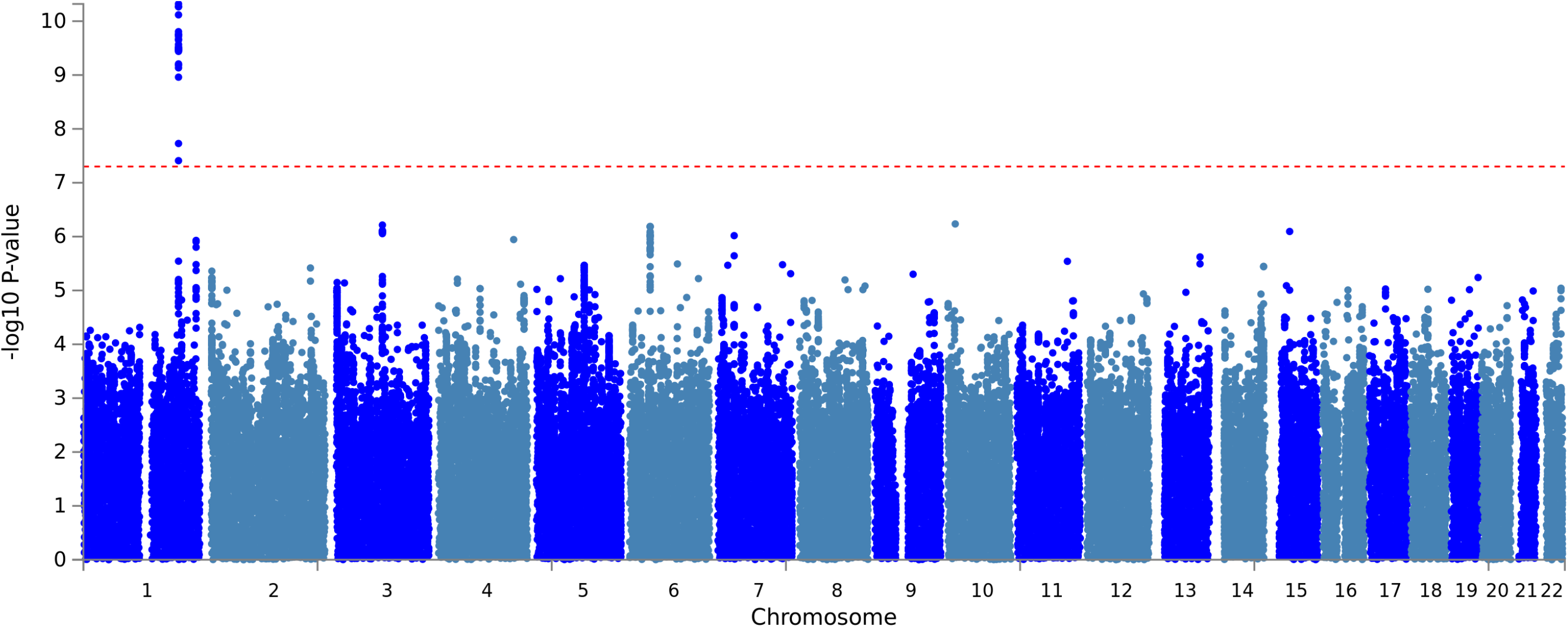
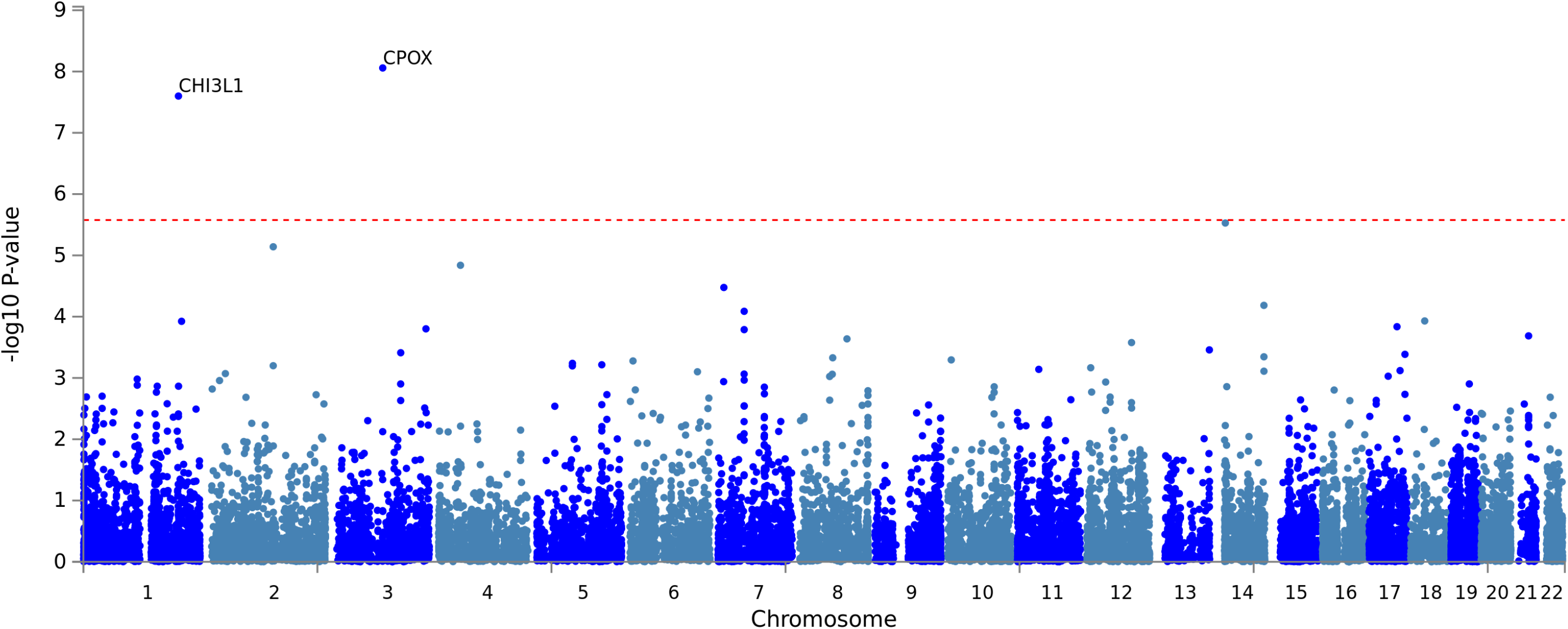
Manhattan plots of **2A)** SNP level and **2B)** gene-level genome-wide association results using log-transformed CSF YKL-40 levels as outcome trait (n = 677). Gene assignments are according to FUMA ^24^. Dotted red lines represent the threshold for genome-wide significance, i.e., α = 5.0E-08 for SNP-based (2A) and α = 2.651E-6 for gene-based (2B) analyses (see Methods).

Gene-based GWAS analyses confirmed the association with *CHI3L1* (P=2.52E-08; Figure 2B, Supplementary Table 4), and revealed a second, independent locus, *CPOX* (coproporphyrinogen oxidase; chromosome 3q11.2-12.1), showing even more significant gene-based association with CSF YKL-40 levels (P = 8.75E-09; Figure 2B, Supplementary Table 4). The most significantly associated single variant in *CPOX* was rs58943879 (P = 6.1E-07; Supplementary Table 4). Manual lookup on the GTEx portal (v8) revealed no previously observed eQTLs in brain, despite CPOX’s relatively pronounced expression in all brain tissues sampled in GTEx (URL: https://www.gtexportal.org/home/gene/CPOX).

### GWAS analyses using CSF neurogranin (Ng) levels

Lastly, the GWAS using CSF Ng levels yielded no genome-wide significant association in the SNP-based analyses (Figure 3A, Table 2, and Supplementary Table 7). The top-ranking SNP-based finding was elicited by rs10052776 (P = 1.0E-07, Supplementary Table 7), located in *CTNND2* mapping to chromosome 5p15.2. Interestingly, SNPs in this gene were previously associated with both late-onset AD and cognitive performance by GWAS ^30,31^ according to the “GWAS catalog” (https://www.ebi.ac.uk/gwas/genes/CTNND2). Gene-based association analyses using MAGMA also did not reveal any genome-wide significant signals with variants annotated to the 18,862 genes utilized in these analyses (Figure 3B). The top-ranking gene-based finding with 21 SNPs was observed with *KDELR1* (P=2.29E-05) mapping to chromosome 19q13.33, a gene hitherto not associated with the traits listed in the “GWAS catalog”.

**Figure 3.**
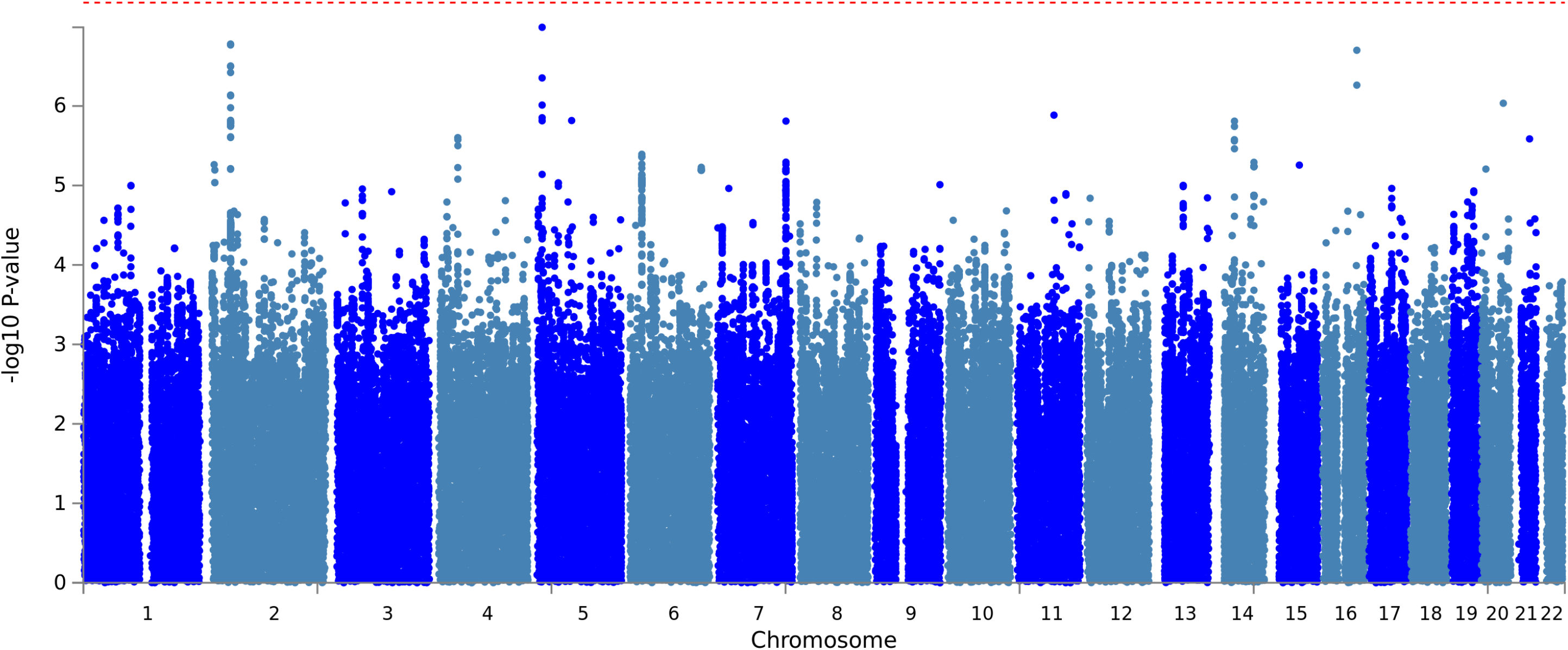
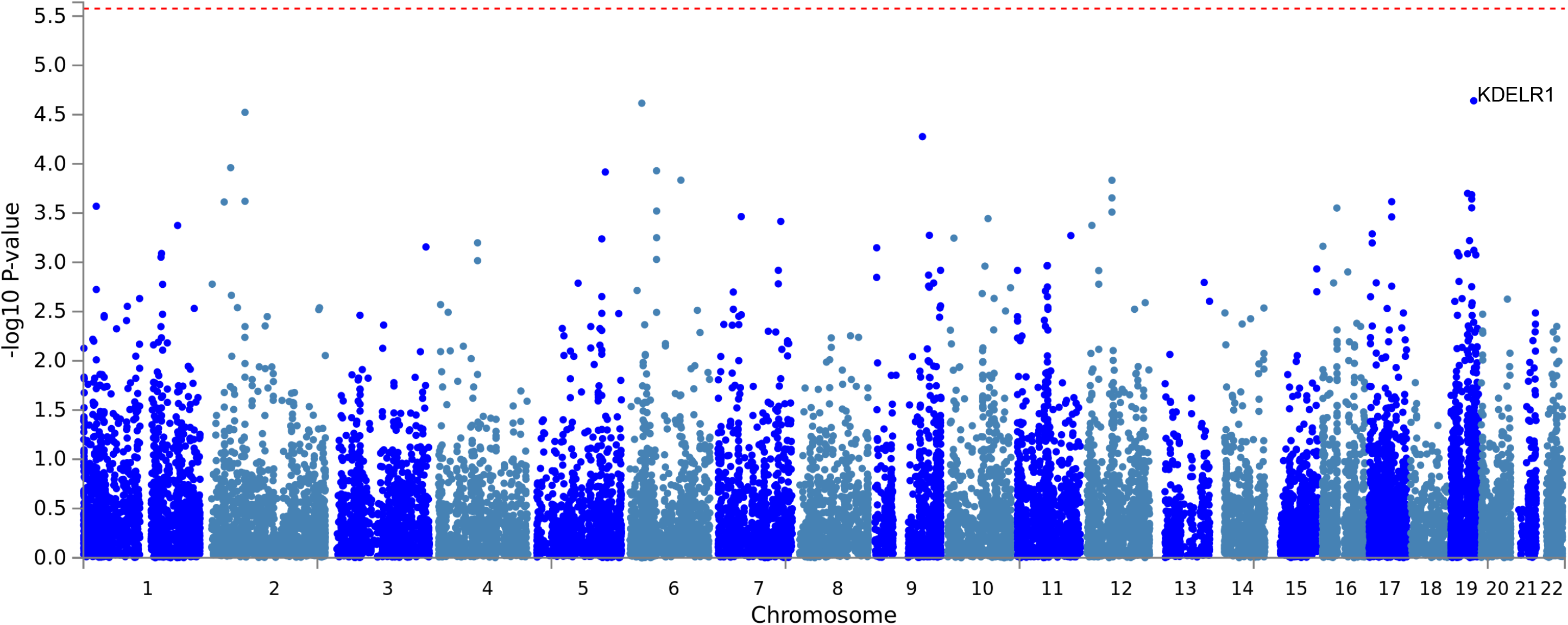
Manhattan plots of **3A)** SNP level and **3B)** gene-level genome-wide association results using log-transformed CSF Ng levels as outcome trait (n = 672). Gene assignments are according to FUMA 24. Dotted red lines represent the threshold for genome-wide significance, i.e., α = 5.0E-08 for SNP-based (3A) and α = 2.651E-6 for gene-based (3B) analyses (see Methods).

### Genetic correlation analyses

To assess whether and to which degree variation of the three AD CSF biomarkers analyzed in this study show association with “AD-related” phenotypes (using summary statistics from two previously reported large AD case-control GWAS ^1,2^), we initially pursued three different analysis paradigms: i) linkage disequilibrium (LD) score regression as implemented in LDSC software ^32^, ii) genetic correlation analyses as implemented in GCTA ^33^, and iii) polygenic risk score (PRS) analysis (see methods). Likely due to the comparatively small sample size of our CSF GWAS dataset, neither LDSC (recommended n>>5,000) nor GCTA-based (recommended n>3,500) yielded interpretable results. In contrast, application of AD-risk GWAS derived PRS as predictors of CSF biomarker variation in approach iii) did yield interpretable associations. However, for both datasets the CSF biomarker phenotype variance in EMIF-AD MDB explained by AD-PRS was collectively minor, reaching nominal significance for some CSF phenotypes and P-value thresholds using Kunkle et al.^2^, but none from the Jansen et al. ^1^ data (Supplementary Table 8). This is in contrast to applying AD GWAS-based PRS to Aβ-related CSF phenotypes in the same EMIF-AD MDB dataset: here, the strongest associations explained nearly 6% of the phenotypic variance (P = 9E-09) ^17^, suggesting that the genetic architectures underlying AD risk and variation at CSF NF-L, YKL-40, and Ng in general do not show any substantial overlap.

## Discussion

We performed GWAS analyses for three CSF AD-related biomarkers in the EMIF-AD MDB dataset and identified novel genome-wide significant association with genetic markers in the established FTLD risk gene *TMEM106B* and NF-L protein concentrations in the CSF. Additional novel signals were observed with DNA variants in *CPOX* and CSF levels of YKL-40. Finally, we detected very strong and genome-wide significant association between markers in *CHI3L1* and CSF levels of YKL-40, representing the only *cis* pQTL finding in our analyses. To the best of our knowledge, our study is the first *bona fide* GWAS for all three of these CSF biomarkers with the exception of two small (n = 133 and n = 265) CSF pQTL GWAS on YKL-40 in people of Asian descent ^34^ and a GWAS on NF-L in non-demented elderly from the the Alzheimer Disease Neuroimaging (ADNI) cohort ^35^. Of note, the former study also identified strong and genome-wide significant *cis* pQTL effects at the *CHI3L1* / YKL-40 locus, corroborating our findings. Other noteworthy results from our study include evidence for several rare-variant associations with CSF NF-L levels and an overall lack of AD-related genetic association signals with the CSF biomarkers analyzed here. This latter point explicitly includes genetic variants in or near the apolipoprotein E gene (*APOE*), the strongest currently known genetic AD risk factor ^3^, which did not show association with any of the CSF biomarkers analyzed here.

Possibly the most noteworthy novel signal observed in our study relates to the association between *TMEM106B* and CSF NF-L. DNA variants in *TMEM106B* have first been implicated in neuropsychiatric research by a GWAS on FTLD with TAR DNA-binding protein (TDP-43) inclusions (FTLD-TDP; Van Deerlin VM et al. ^27^), a finding that was subsequently confirmed in independent datasets (see Pottier C et al. ^28^ for recent GWAS results). Furthermore, a recent meta-analysis revealed that CSF NF-L levels are significantly increased in FTLD ^8^. In addition to FTLD, the “GWAS catalog” database lists a number of other, mostly neuropsychiatric (e.g., depression, differential brain aging, neuroticism) but also non-neurological (e.g., leukemia, height, HDL levels) traits showing genomewide association with *TMEM106B* (URL: https://www.ebi.ac.uk/gwas/genes/TMEM106B; Buniello A et al. ^36^). Furthermore, there is a strong *cis* pQTL GWAS evidence for SNPs in this region (in particular: rs10950398) and *TMEM106B* protein levels in blood ^37^. In GTEx, this same SNP is also reported to correlate with *TMEM106B* mRNA expression in brain (cortex and cerebellum), albeit at lesser significance (https://www.gtexportal.org/home/snp/rs10950398). Of note, this pQTL / eQTL variant (rs10950398) is in nearly perfect LD with the lead SNP identified here to show association with CSF NF-L levels (i.e., rs1548884, r^2^ = 0.97; Supplementary Tables 1 & 2). In summary, there is now convincing evidence converging from multiple lines of independent data that DNA variants in *TMEM106B* not only show association with several neuropsychiatric and non-neurological phenotypes, but also *cis* (*TMEM106B*, previous work) and *trans* (CSF NF-L, this study) pQTL associations with proteins relevant for neuronal function. In the AD field, NF-L recently (re)gained interest based on data suggesting that NF-L protein dynamics in serum may predict progression and brain neurodegeneration at early pre-symptomatic stages of familial AD ^9^, and tracks neurodegeneration in sporadic AD ^38^. The novel results from our GWAS indicate that DNA sequence variants in *TMEM106B* may be involved in regulating CSF NF-L protein levels. Since the same variant(s) are also *cis* eQTLs / pQTLs of *TMEM106B* mRNA / protein levels, it is tempting to speculate that the observed effect on CSF NF-L may be mediated by *TMEM106B* mRNA or protein. In line with this hypothesis is the observation that the lead genetic variant in *TMEM106B* highlighted in our analyses (rs1548884) shows some, albeit sub genome-wide, evidence for association with AD risk in the two largest and most recent GWAS in the field, i.e., P = 0.00005 and P=0.005 in Jansen et al. ^1^ and Kunkle et al. ^2^, respectively. In conclusion, our novel data now provide the genetic foundation for future work aimed at elucidating whether the observed increase in CSF NF-L levels represents a “cause” or “effect” of the neurodegenerative processes underlying symptomatic and pre-symptomatic AD. The observation that the same *TMEM106B* variants show association with both AD risk *and* CSF NF-L levels in independent datasets provide a first indication that the recently proposed change in “NF-L dynamics” ^9^ may, indeed, contribute to AD neuropathology rather than simply reflecting an effect of the same. However, additional work is needed, e.g. replication of the original NF-L dynamics result and approaches applying Mendelian Randomization, to address this question more formally. Interestingly, across the neurodegenerative dementias, FTLD is among the diseases with the highest CSF NF-L concentrations ^8^.

To our knowledge, the only other GWAS investigating CSF NF-L was performed on 265 non-demented individuals from the ADNI cohort ^35^. These authors highlighted two SNPs (i.e. rs465401 and rs460420) in *ADAMTS1* to show genome-wide significant association with CSF NF-L. However, these results were not replicated in the EMIF-AD MBD dataset, where these SNPs showed P-values of 0.5485, and 0.5391, respectively. Conversely, variants in *TMEM106B* were not highlighted as “peak results” in that study. However, apart from the difference in sample size, there were several additional noteworthy differences in the analysis approach used by us and Niu et al. ^35^ (e.g. they did not use genotype imputations to increase their coverage of untyped portions of the genome; in addition to sex and genetic ancestry, they also included age, education years, and *APOE* e4 status as covariates in the primary GWAS analyses). Future work in more individuals needs to determine whether the association between *TMEM106B* and CSF NF-L observed here will prove genuine.

The second novel result emerging from our analyses imply that variants in *CPOX* (encoding coproporphyrinogen oxidase) are associated with YKL-40 levels in CSF. *CPOX* is ubiquitously expressed (based on GTEx release [v8]) and encodes an enzyme involved in the heme biosynthetic pathway. Intracellularly, it localizes to the mitochondria and catalyzes the 2-step oxidative decarboxylation of the heme precursor coproporphyrinogen III to protoporphyrinogen IX (https://omim.org/entry/612732). While common variants in this gene have hitherto not been associated with any human trait recorded in the “GWAS catalog”, rare mutations in *CPOX* can cause coproporphyria and harderoporphyria (OMIM phenotype ID # 121300), hereditary forms of porphyrias characterized by enzyme deficiencies in the heme biosynthetic pathway. Previous work has suggested that heme has a strong affinity for binding Aβ-42 peptide *in vitro*^39^,, leading to speculations that porphyrias – through mechanisms involved in the inherent intracellular heme deficiency caused by these diseases – could potentially alter the risk and / or course of AD ^40^. However, these potential links have not hitherto been directly proven in experimental or other work and must be considered “speculative”. Likewise, it remains unclear how variation in *CPOX* mRNA or protein expression should impact levels of YKL-40 in the CSF.

Finally, the third main finding worth discussing is the *cis* pQTL result linking markers in *CHI3L1* to CSF levels of YKL-40, which represents the strongest and most significant of all association signals in our GWAS. Several prior publications have implicated genetic variants in *CHI3L1* to represent *cis* pQTLs of YKL-40 levels in blood ^37,41–43^ (for more details see: https://www.ebi.ac.uk/gwas/genes/CHI3L1). However, to the best of our knowledge, only one prior study has investigated YKL-40 levels in the human CSF ^34^. Notwithstanding that study’s relatively small sample size (n=133) and different ethnicity (Japanese), this GWAS, too, reported very pronounced *cis* pQTL effects of genetic variants in *CHI3L1*. Taken together, there is now compelling converging evidence that expression of YKL-40 in both blood and CSF is regulated by DNA sequence variants located in the very gene encoding this protein. These same variants are also found as eQTL and mQTLs in independent datasets. However, unlike the situation observed for our GWAS results for NFL, the YKL-40 regulatory SNPs do not show any evidence of association with AD risk in the GWAS by Jansen et al. ^1^ and Kunkle et al. ^2^. Thus, owing to this general absence of genetic association with AD risk it appears that the observed association between CSF YKL-40 and AD status ^7^ probably lies downstream of the initiation of AD neuropathology.

While our data highlights some important new insights into the genetic architecture underlying three CSF AD biomarkers and, hence, into the mechanisms underlying AD neuropathogenesis, our study is subject to some limitations. First and foremost, owing to the lack of appropriate datasets with both genome-wide SNP genotype and CSF data of the three biomarkers in question, we were unable to independently validate our association signals. For several other variables collected in EMIF-AD MDB, data from ADNI project has served as valuable independent reference, including some analyzed by GWAS (e.g. Hong S et al. ^17^). While ADNI investigators are in the process of collecting CSF data for NF-L, YKL-40, and Ng, data available with genetic information at the time of this study only ranged from n=82 to 125 in the whole-genome sequencing (WGS) subset of ADNI, precluding any meaningful GWAS analysis in this setting. Second, although our dataset is the first and / or largest to allow GWAS analyses on all three CSF variables covered, the sample size available for analysis is still relatively modest (range: 671 to 677), limiting our power to detect genetic variants of moderate to small effects. Thus, the results of our GWAS likely only represent the “lowest hanging fruit” of the genetic factors underlying the analyzed traits. Third, we note that while both the genome-wide SNP genotypes as well as CSF biomarker concentrations were generated in one run of consecutive experiments in two dedicated laboratories (one for genotyping, one for CSF markers; likely reducing the possibility of batch effects), the CSF specimen were collected individually at each of the 11 EMIF-AD MDB participating sites, sometimes using different collection procedures and CSF storage tubes. While this sampling heterogeneity could have affected all or some of our results (although YKL-40 and Ng were recently shown to be quite stable across a range of conditions ^44^, we note that CSF drawing was performed independent of genotype, so any batch effect in this particular setting should be minimal if present at all. Finally, we note that the EMIF-AD MBD dataset was not designed to be “representative” of the general population but was assembled with the aim to achieve approximately equal proportions of “amyloid+” vs. “amyloid-“ individuals across in individuals with normal cognition and MCI although this was only achieved for MCI (see Methods and Bos I et al. ^16^). While this ascertainment strategy does not invalidate our GWAS results, they may not be generalizable to the population as a whole. However, this limitation can affect any study with clinically ascertained participants and, thus, applies to most previously published GWAS in the field, including those performed in ADNI.

In conclusion, our GWAS on CSF NF-L, YKL-40 and Ng levels provides important new insights into the genetic architecture underlying inter-individual variation in these traits. Together with recent GWAS results on AD risk from case-control studies, they shed important new light on the sequence of events in relation to the initiation and progression of neuropathological processes relevant in AD. Additional work is needed to set our results onto a broader evidence-based foundation and to clarify the molecular mechanisms underlying the observed associations.

## Author contributions

SH performed all the analyses and interpretation on data and co-wrote all drafts of the manuscript. VD extracted EMIF-AD MBD DNA sample. IB, SV, BT, and PJ coordinated the collection and harmonization of phenotypes and biosamples in EMIF-AD MBD and helped identifying equivalent phenotypes from the ADNI catalog. DP and RT contributed to replication analyses in ADNI. AF and MW supervised the genotyping experiments. UA, KB and HZ performed CSF biomarker measurements and took part in cut-point determinations. KS and CVB contributed to genetic characterization of samples and design of the genomics studies in EMIF-AD MBD. RV and SG, contributed to sample and data collection. IS were responsible for data collection of the Antwerp cohort. JS, PJV and SL are leads for the EMIF-AD MBD; as such they designed and managed the platform. LB designed and supervised the genomics portion of the EMIF-AD MBD project and co-wrote all drafts of the manuscript. All authors critically revised all manuscripts drafts, read and approved the final manuscript.

## Acknowledgements

The present study was conducted as part of the EMIF-AD MBD project, which has received support from the Innovative Medicines Initiative Joint Undertaking under EMIF grant agreement n° 115372, the resources of which are composed of financial contribution from the European Union’s Seventh Framework Program (FP7/2007–2013) and EFPIA companies’ in kind contribution. Parts of this study were made possible through support from the German Research Foundation (DFG grant FOR2488: Main support by subproject “INF-GDAC” BE2287/7-1 to LB) and the Cure Alzheimer’s Fund (to L.B. and R.E.T.). RV acknowledges the support by the Stichting Alzheimer Onderzoek (#13007, #11020, #2017-032) and the Flemish Government (VIND IWT 135043). KB is supported by the Swedish Research Council (#2017-00915), the Alzheimer Drug Discovery Foundation (ADDF), USA (#RDAPB-201809-2016615), the Swedish Alzheimer Foundation (#AF-742881), Hjärnfonden, Sweden (#FO2017-0243), the Swedish state under the agreement between the Swedish government and the County Councils, the ALF-agreement (#ALFGBG-715986), and European Union Joint Program for Neurodegenerative Disorders (JPND2019-466-236). HZ is a Wallenberg Scholar supported by grants from the Swedish Research Council (#2018-02532), the European Research Council (#681712) and Swedish State Support for Clinical Research (#ALFGBG-720931). SJBV received funding from the Innovative Medicines Initiative 2 Joint Undertaking under ROADMAP grant agreement No. 116020 and from ZonMw during the conduct of this study. No conflict of interest exists. Research at VIB-UAntwerp was in part supported by the University of Antwerp Research Fund. The authors acknowledge the assistance of Ellen De Roeck, Naomi De Roeck and Hanne Struyfs (UAntwerp) with data collection. The Lausanne study was funded by a grant from the Swiss National Research Foundation (SNF 320030_141179) to JP.

We thank Mrs. Tanja Wesse and Mrs. Sanaz Sedghpour Sabet at the Institute of Clinical Molecular Biology, Christian-Albrechts-University of Kiel, Kiel, Germany for technical assistance with the GSA genotyping. We thank Dr. Fabian Kilpert for his assistance with the QC and genotype imputations. The LIGA team acknowledges computational support from the OMICS compute cluster at the University of Lübeck.

## Additional files

*The supplementary material supplied with this manuscript contains Supplementary Methods, Supplementary Figures 1-4 and Supplementary Tables 1-8*.

## References

1 Jansen IE, Savage JE, Watanabe K, et al. Genome-wide meta-analysis identifies new loci and functional pathways influencing Alzheimer’s disease risk. Nat Genet 2019; : 1.

2 Kunkle BW, Grenier-Boley B, Sims R, et al. Genetic meta-analysis of diagnosed Alzheimer’s disease identifies new risk loci and implicates Aβ, tau, immunity and lipid processing. Nat Genet 2019; 51: 414–30.

3 Bertram, Lars; Tanzi RE. Genomic Mechanisms in Alzheimer’s Disease. Brain Pathol.

4 Bos I, Vos S, Verhey F, et al. Cerebrospinal fluid biomarkers of neurodegeneration, synaptic integrity, and astroglial activation across the clinical Alzheimer’s disease spectrum. Alzheimer’s Dement 2019; 15: 644–54.

5 Yuan A, Nixon RA. Specialized roles of neurofilament proteins in synapses: Relevance to neuropsychiatric disorders. Brain Res Bull 2016; 126: 334–46.

6 Yuan A, Rao M V., Veeranna, Nixon RA. Neurofilaments at a glance. J. Cell Sci. 2012; 125: 3257–63.

7 Olsson B, Lautner R, Andreasson U, et al. CSF and blood biomarkers for the diagnosis of Alzheimer’s disease: a systematic review and meta-analysis. Lancet Neurol 2016; 15: 673–84.

8 Bridel C, Van Wieringen WN, Zetterberg H, et al. Diagnostic Value of Cerebrospinal Fluid Neurofilament Light Protein in Neurology: A Systematic Review and Meta-analysis. JAMA Neurol 2019; 76: 1035–48.

9 Preische O, Schultz SA, Apel A, et al. Serum neurofilament dynamics predicts neurodegeneration and clinical progression in presymptomatic Alzheimer’s disease. Nat. Med. 2019; 25: 277–83.

10 Molinuevo JL, Ayton S, Batrla R, et al. Current state of Alzheimer’s fluid biomarkers. Acta Neuropathol 2018; 136: 821–53.

11 Querol-Vilaseca M, Colom-Cadena M, Pegueroles J, et al. YKL-40 (Chitinase 3-like I) is expressed in a subset of astrocytes in Alzheimer’s disease and other tauopathies. J Neuroinflammation 2017; 14. DOI:10.1186/s12974-017-0893-7.

12 Chang JW, Schumacher E, Coulter PM, Vinters H V, Watson JB. Dendritic translocation of RC3/neurogranin mRNA in normal aging, Alzheimer disease and fronto-temporal dementia. J Neuropathol Exp Neurol 1997; 56: 1105–18.

13 Scheff SW, Price DA, Schmitt FA, DeKosky ST, Mufson EJ. Synaptic alterations in CA1 in mild Alzheimer disease and mild cognitive impairment. Neurology 2007; 68: 1501–8.

14 Slemmon JR, Feng B, Erhardt JA. Small Proteins that Modulate Calmodulin-Dependent Signal Transduction. Mol Neurobiol 2000; 22: 099–114.

15 Tarawneh R, D’Angelo G, Crimmins D, et al. Diagnostic and prognostic utility of the synaptic marker neurogranin in Alzheimer disease. JAMA Neurol 2016; 73: 561–71.

16 Bos I, Vos S, Vandenberghe R, et al. The EMIF-AD Multimodal Biomarker Discovery study: design, methods and cohort characteristics. Alzheimers Res Ther 2018; 10: 64.

17 Hong S, Prokopenko D, Dobricic V, et al. Genome-wide association study of Alzheimer’s disease CSF biomarkers in the EMIF-AD Multimodal Biomarker Discovery dataset. BioRxiv 2019; : 774554.

18 Zetterberg H, Skillbäck T, Mattsson N, et al. Association of cerebrospinal fluid neurofilament light concentration with Alzheimer disease progression. JAMA Neurol 2016. DOI:10.1001/jamaneurol.2015.3037.

19 Portelius E, Zetterberg H, Skillbäck T, et al. Cerebrospinal fluid neurogranin: Relation to cognition and neurodegeneration in Alzheimer’s disease. Brain 2015. DOI:10.1093/brain/awv267.

20 Olsson B, Hertze J, Lautner R, et al. Microglial markers are elevated in the prodromal phase of Alzheimer’s disease and vascular dementia. J Alzheimer’s Dis 2013. DOI:10.3233/JAD-2012-120787.

21 Li Y, Willer CJ, Ding J, Scheet P, Abecasis GR. MaCH: using sequence and genotype data to estimate haplotypes and unobserved genotypes. Genet Epidemiol 2010; 34: 816–34.

22 Li Y, Willer C, Sanna S, Abecasis G. Genotype Imputation. Annu Rev Genomics Hum Genet 2009; 10: 387–406.

23 Aulchenko YS, Ripke S, Isaacs A, van Duijn CM. GenABEL: an R library for genome-wide association analysis. Bioinformatics 2007; 23: 1294–6.

24 Watanabe K, Taskesen E, van Bochoven A, Posthuma D. Functional mapping and annotation of genetic associations with FUMA. Nat Commun 2017; 8: 1826.

25 de Leeuw CA, Mooij JM, Heskes T, Posthuma D. MAGMA: Generalized Gene-Set Analysis of GWAS Data. PLOS Comput Biol 2015; 11: e1004219.

26 Purcell S, Neale B, Todd-Brown K, et al. PLINK: a tool set for whole-genome association and population-based linkage analyses. Am J Hum Genet 2007; 81: 559–75.

27 Van Deerlin VM, Sleiman PMA, Martinez-Lage M, et al. Common variants at 7p21 are associated with frontotemporal lobar degeneration with TDP-43 inclusions. Nat Genet 2010; 42: 234–9.

28 Pottier C, Zhou X, Perkerson RB, et al. Potential genetic modifiers of disease risk and age at onset in patients with frontotemporal lobar degeneration and GRN mutations: a genome-wide association study. Lancet Neurol 2018; 17: 548–58.

29 Gaunt TR, Shihab HA, Hemani G, et al. Systematic identification of genetic influences on methylation across the human life course. Genome Biol 2016; 17: 61.

30 Lee JJ, Wedow R, Okbay A, et al. Gene discovery and polygenic prediction from a genomewide association study of educational attainment in 1.1 million individuals. Nat Genet 2018; 50: 1112–21.

31 Mez J, Chung J, Jun G, et al. Two novel loci, COBL and SLC10A2, for Alzheimer’s disease in African Americans. Alzheimer’s Dement 2017; 13: 119–29.

32 Bulik-Sullivan B, Finucane HK, Anttila V, et al. An atlas of genetic correlations across human diseases and traits. Nat Genet 2015; 47: 1236–41.

33 Byers SA, Price JP, Cooper JJ, Li Q, Price DH. HEXIM2, a HEXIM1-related protein, regulates positive transcription elongation factor b through association with 7SK. J Biol Chem 2005; 280: 16360–7.

34 Sasayama D, Hattori K, Ogawa S, et al. Genome-wide quantitative trait loci mapping of the human cerebrospinal fluid proteome. Hum Mol Genet 2017. DOI:10.1093/hmg/ddw366.

35 Niu L-D, Xu W, Li J-Q, et al. Genome-wide association study of cerebrospinal fluid neurofilament light levels in non-demented elders. Ann Transl Med 2019; 7: 657–657.

36 Buniello A, Macarthur JAL, Cerezo M, et al. The NHGRI-EBI GWAS Catalog of published genome-wide association studies, targeted arrays and summary statistics 2019. Nucleic Acids Res 2019. DOI:10.1093/nar/gky1120.

37 Emilsson V, Ilkov M, Lamb JR, et al. Co-regulatory networks of human serum proteins link genetics to disease. Science (80-) 2018. DOI:10.1126/science.aaq1327.

38 Mattsson N, Cullen NC, Andreasson U, Zetterberg H, Blennow K. Association Between Longitudinal Plasma Neurofilament Light and Neurodegeneration in Patients With Alzheimer Disease. JAMA Neurol 2019; 76: 791.

39 Atamna H, Frey WH. A role for heme in Alzheimer’s disease: Heme binds amyloid β and has altered metabolism. Proc Natl Acad Sci U S A 2004. DOI:10.1073/pnas.0404349101.

40 Dwyer BE, Stone M, Zhu X, Perry G, Smith MA. Heme deficiency in Alzheimer’s disease: A possible connection to porphyria. J. Biomed. Biotechnol. 2006; 2006. DOI:10.1155/JBB/2006/24038.

41 Sun BB, Maranville JC, Peters JE, et al. Genomic atlas of the human plasma proteome. Nature 2018. DOI:10.1038/s41586-018-0175-2.

42 Suhre K, Arnold M, Bhagwat AM, et al. Connecting genetic risk to disease end points through the human blood plasma proteome. Nat Commun 2017. DOI:10.1038/ncomms14357.

43 Ober C, Tan Z, Sun Y, et al. Effect of variation in CHI3L1 on serum YKL-40 level, risk of asthma, and lung function. N Engl J Med 2008. DOI:10.1056/NEJMoa0708801.

44 Willemse EAJ, Vermeiren Y, Garcia-Ayllon MS, et al. Pre-analytical stability of novel cerebrospinal fluid biomarkers. Clin Chim Acta 2019; 497: 204–11.

