## Supplementary Material for "*TMEM106B* and *CPOX* are genetic determinants of cerebrospinal fluid Alzheimer’s disease biomarker levels"

### Supplementary Methods

*NB: Some text in the methods description of this document was taken from our previous EMIF-AD MDB GWAS manuscript Hong et al. [1] without explicit referencing.*

#### Sample description

The ascertainment procedures for the EMIF-AD cohort are described elsewhere [2]. In brief, the dataset includes 1221 elderly individuals (years of age: mean = 67.9, SD = 8.3) with different cognitive diagnoses at baseline (NC = normal cognition; MCI = mild cognitive impairment; AD = AD-type dementia) and the clinical follow-up data were available for 759 individuals. Depending on the availability of the clinical records, each phenotype has different effective sample sizes (Table 1). The demographic information for the three quantitative CSF phenotypes of the EMIF-AD MBD dataset utilized in this paper is summarized in Table 1.

#### CSF biomarkers

Details of the CSF biomarker measurements can be found in Bos et al. [3]. In brief, CSF samples were shipped to Department of Psychiatry and Neurochemistry at University of Gothenburg, Sweden. NF-L levels were measured using a commercial ELISA (NF-light ELISA, UmanDiagnostics, Umea, Sweden [4]). Ng levels were measured using an in-house immunoassay for Ng [5]. YKL-40 levels were measured using a human chitinase-3 quantikine ELISA kit (R&D systems, Inc, Minneapolis, MN [6]). All analyses were performed according to the manufacturer's instructions. To reduce the skewness of the phenotype distributions, all three CSF outcome variables utilized in this study were log-transformed prior to analysis (see Supplementary figure 1 for density plots pre- and post-transformation).

**Genotype data handling, QC and imputation procedures**

Genotyping was performed at the UKSH NGS facility located at the Institute for Clinical Molecular Biology (IKMB) located in Kiel, Germany. All post-genotyping data processing and handling was performed at LIGA located at UKSH campus Lübeck / University of Lübeck.

Raw data processing, i.e. clustering and genotype calling from raw intensity data (idat format) was performed in GenomeStudio software (v2.0.4; Illumina, Inc.) using the genotyping module (version 2.0.2). Samples with call rate <0.95 and p50GC <0.7 were excluded at this stage. We then used PLINK software **v1.9** [7] to perform additional QC filtering, i.e. sex checks (--check-sex 0.25 0.75), strand check (--flip), missing genotype rate (--geno 0.02; --mind 0.05), Hardy-Weinberg equilibrium (HWE) tests (--hwe 0.000005),   and minor allele frequency (MAF) filtering (--maf 0.01). For determining pairwise allele sharing (to identify cryptic relatedness), we used an LD-pruned set of markers (--indep-pairwise 1500 150 0.2).   Pairwise allele-sharing IBD/IBS was determined using (--Z-genome --min 0.1). Overall, this procedure led to 498,589 QC-filtered SNPs in 931 samples suitable for imputation. The LD pruned dataset was also used for principal component analysis (PCA; using PLINK command "--pca") along with the reference dataset of the 1000 Genome Project Consortium Phase 3 (The 1000 Genomes Project Consortium, 2015) to assign ethnic descent groups using the five 1000G super-populations by k-nearest neighbor (k-NN; k=9) classification (using R package 'class' [8] in R 2.3.2). This resulted in assigning a "European descent" to 898 out of all 931 samples; only these n=898 samples were used in the subsequent statistical analyses.

Before imputation, the QC'ed genotype data were then subjected to bcftools (v1.9) [9] for removing ambiguous SNPs, and flipping and swapping alleles to align to GRCh37/hg19. This was followed by haplotype phasing using SHAPEIT2 [10] and imputation of unobserved genotypes using Minimac3 [11] using a precompiled Haplotype Reference Consortium (HRC) reference panel (EGAD00001002729 including 39,131,578 SNPs from ~11K individuals). Post-imputation we only retained autosomal SNPs with minimac3 Rsq ≥0.3, MAF ≥1%, and HWE P-values ≥ 5E-6 (using best-guess genotypes in control subjects), leaving a total of 7,778,465 SNPs for statistical analyses.

**GWAS and post-GWAS analyses**

Linear regression was utilized to perform SNP-based association analyses using imputation-derived allele dosages as independent variables and the log-transformed concentrations of CSF NF-L, CSF Ng, and CSF YKL-40 as dependent outcome variables (utilizing mach2qtl software [12,13]). Covariates included into the regression models were sex, age at examination, diagnostic groups (coded as AD = 3, MCI = 2, controls = 1) as well as ancestry-specific principal components (PCs) 1-5 derived from PC analysis on a linkage-disequilibrium pruned set of SNPs using PLINK v1.9 [7]. The genomic inflation factor for both SNP- and gene-based analyses was calculated in R using the "GenABEL" package and is depicted on the QQ plots in the Supplementary Material [14]. Genome-wide significance for the SNP-based analyses was defined as α = 5E-08, a widely used threshold that accounts for the approximate number of independent variants (~1M) in European populations [15,16]. We used FUMA (http://fuma.ctglab.nl/) [17] for post-GWAS functional mapping and annotation of the genome-wide association. This included performing gene-based GWAS analyses using MAGMA [18] as implemented in FUMA. As suggested on the FUMA website, the threshold of genome-wide significance in the gene-based analyses was defined as α=0.05/18720 = 2.671E-06 based on the number of genes (n=18720) utilized for these analyses.

**Polygenic risk score (PRS) analysis**

To aggregate data on multiple variants per individual we computed polygenic risk scores (PRS) for each individual which were then used as independent variable in the statistical analyses. Allele status and effect-size estimates were taken from the summary statistics of the two largest AD GWAS published to date, i.e. the Jansen et al. [19], and Kunkle et al. [20] publications. First, we removed ambiguous SNPs (A/T and C/G) from the list of considered variants. Furthermore, we only used SNPs with MAF >0.01 and imputation quality $r^2$ >0.8 in the EMIF-AD dataset. Next, LD pruning was performed on the CEU portion of the HRC reference panel used for imputation. To this end, we used PLINK 1.9 software [7] for two consecutive rounds of marker pruning (1st round: --clump-p1 1 --clump-p2 1 --clump-r2 .5 --clump-kb 250; 2nd round: --clump-p1 1 --clump-p2 1 --clump-r2 .2 --clump-kb 5000). Actual PRS were computed using the --score command in PLINK for a variety of P-value thresholds in the primary GWAS data (i.e 5E-08, 5E-06, 1E-04, 0.01, 0.05, 0.10, 0.20, 0.30, 0.40, 0.50, 1.00). The resulting PRS were used as independent variable in the linear or logistic regression models adjusting for sex, age, and PC1 to PC5 as covariates. For the phenotypes not representing the diagnostic outcome, we also included diagnosis as additional covariate. For linear models, variance explained ($R^2$) was derived from comparing results from the full model (including outcome phenotype and covariates) vs the null model (linear model with covariates only). For logistic models, we calculated Nagelkerke's $r^2$ using the R package fmsb.

### Reference

1 Hong S, Prokopenko D, Dobricic V, *et al.* Genome-wide association study of Alzheimer's disease CSF biomarkers in the EMIF-AD Multimodal Biomarker Discovery dataset. *bioRxiv* 2020; : 36.

2 Bos I, Vos S, Vandenberghe R, *et al.* The EMIF-AD Multimodal Biomarker Discovery study: design, methods and cohort characteristics. *Alzheimers Res Ther* 2018; **10**: 64.

3 Bos I, Vos S, Verhey F, *et al.* Cerebrospinal fluid biomarkers of neurodegeneration, synaptic integrity, and astroglial activation across the clinical Alzheimer's disease spectrum. *Alzheimer's Dement* 2019. DOI:10.1016/j.jalz.2019.01.004.

4 Zetterberg H, Skillbäck T, Mattsson N, *et al.* Association of cerebrospinal fluid neurofilament light concentration with Alzheimer disease progression. *JAMA Neurol* 2016. DOI:10.1001/jamaneurol.2015.3037.

5 Portelius E, Zetterberg H, Skillbäck T, *et al.* Cerebrospinal fluid neurogranin: Relation to cognition and neurodegeneration in Alzheimer's disease. *Brain* 2015. DOI:10.1093/brain/awv267.

6 Olsson B, Hertze J, Lautner R, *et al.* Microglial markers are elevated in the prodromal phase of Alzheimer's disease and vascular dementia. *J Alzheimer's Dis* 2013. DOI:10.3233/JAD-2012-120787.

7 Purcell S, Neale B, Todd-Brown K, *et al.* PLINK: A Tool Set for Whole-Genome Association and Population-Based Linkage Analyses. *Am J Hum Genet* 2007. DOI:10.1086/519795.

8 Venables WN, Springer BDR. Modern Applied Statistics with S Fourth edition. http://www.insightful.com. (accessed May 24, 2019).

9 Narasimhan V, Danecek P, Scally A, Xue Y, Tyler-Smith C, Durbin R. BCFtools/RoH: A hidden Markov model approach for detecting autozygosity from next-generation sequencing data. *Bioinformatics* 2016. DOI:10.1093/bioinformatics/btw044.

10 Delaneau O, Marchini J, Zagury JF. A linear complexity phasing method for thousands of genomes. *Nat Methods* 2012. DOI:10.1038/nmeth.1785.

11 Das S, Forer L, Schönherr S, *et al.* Next-generation genotype imputation service and methods. *Nat Genet* 2016. DOI:10.1038/ng.3656.

12 Li Y, Willer C, Sanna S, Abecasis G. Genotype Imputation. *Annu Rev Genomics Hum Genet* 2009; **10**: 387–406.

13 Li Y, Willer CJ, Ding J, Scheet P, Abecasis GR. MaCH: using sequence and genotype data to estimate haplotypes and unobserved genotypes. *Genet Epidemiol* 2010; **34**: 816–34.

14 Aulchenko YS, Ripke S, Isaacs A, van Duijn CM. GenABEL: an R library for genome-wide association analysis. *Bioinformatics* 2007; **23**: 1294–6.

15 International HapMap Consortium TIH. A haplotype map of the human genome. *Nature* 2005; **437**: 1299–320.

16 Pe'er I, Yelensky R, Altshuler D, Daly MJ. Estimation of the multiple testing burden for genomewide association studies of nearly all common variants. *Genet Epidemiol* 2008; **32**: 381–5.

17 Watanabe K, Taskesen E, van Bochoven A, Posthuma D. Functional mapping and annotation of genetic associations with FUMA. *Nat Commun* 2017; **8**: 1826.

18 de Leeuw CA, Mooij JM, Heskes T, Posthuma D. MAGMA: Generalized Gene-Set Analysis of GWAS Data. *PLOS Comput Biol* 2015; **11**: e1004219.

19 Jansen IE, Savage JE, Watanabe K, *et al.* Genome-wide meta-analysis identifies new loci and functional pathways influencing Alzheimer's disease risk. *Nat Genet* 2019; : 1.

20 Kunkle BW, Grenier-Boley B, Sims R, *et al.* Genetic meta-analysis of diagnosed Alzheimer's disease identifies new risk loci and implicates Aβ, tau, immunity and lipid processing. *Nat Genet* 2019; **51**: 414–30.

#### Supplementary Tables

Supplementary Tables 1-8 are can be found in the MS-Excel file "hong.supplementary_tables.xls".

#### Supplementary Figures

Supplementary Figures 2-4 are each arranged in the same format ("X" stands for figure number). See Table 1 in the main text for a full description of variable names.

Fig XA: quantile-to-quantile plots of **SNP-based** (top) and **gene-based** (bottom) genome-wide association results. Lambda values as measures of genomic inflation of the respective analysis are provided alongside the plots.

Fig XB: Regional plots of genome-wide association results drawn in FUMA.

**Figure S1.** Density plot of three CSF biomarkers and their log-transformed value

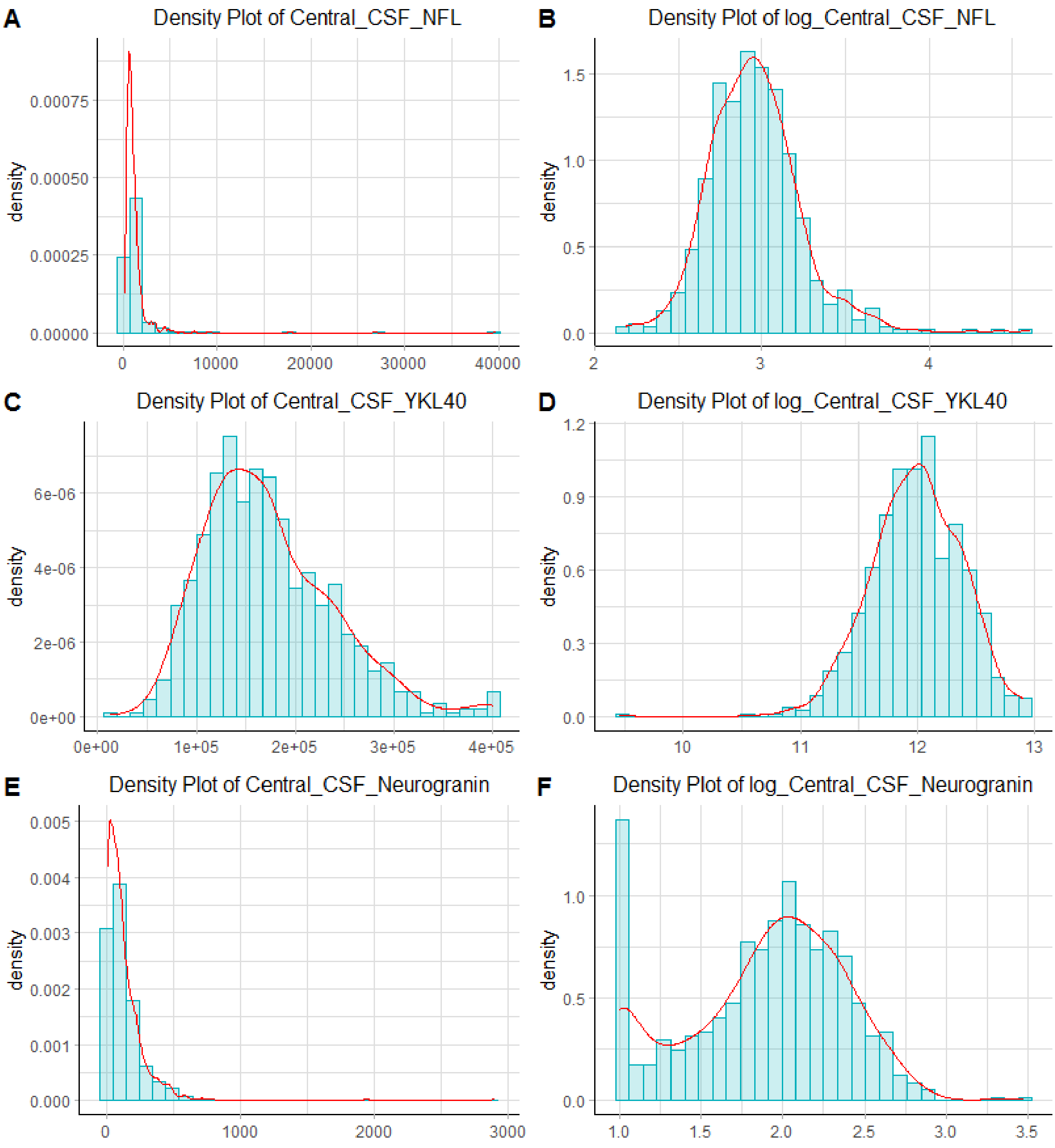

**Figure S2a.** log-transformed CSF_NF-L

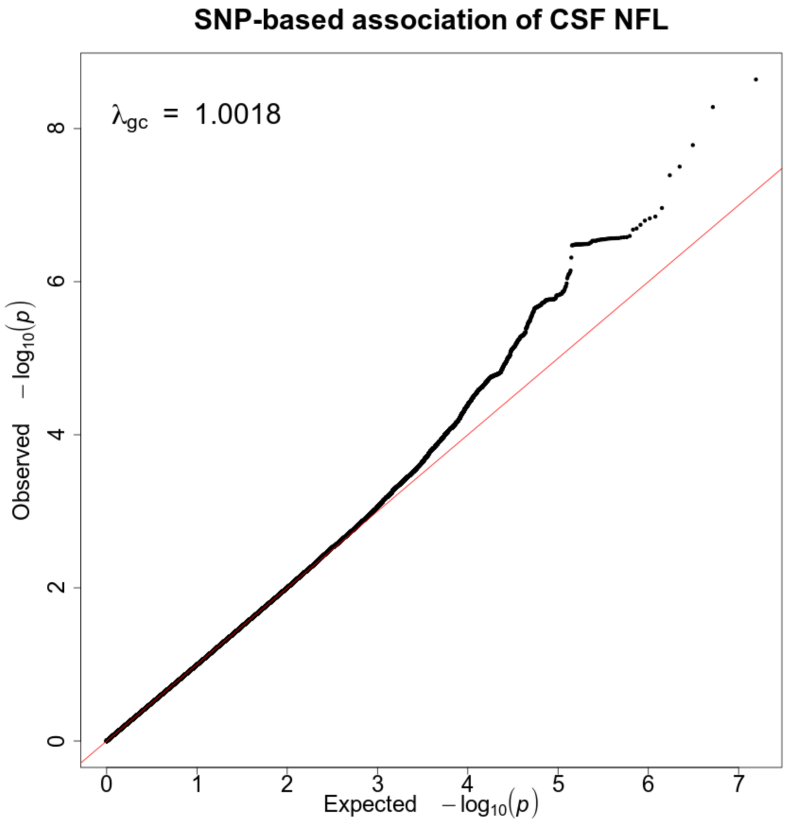


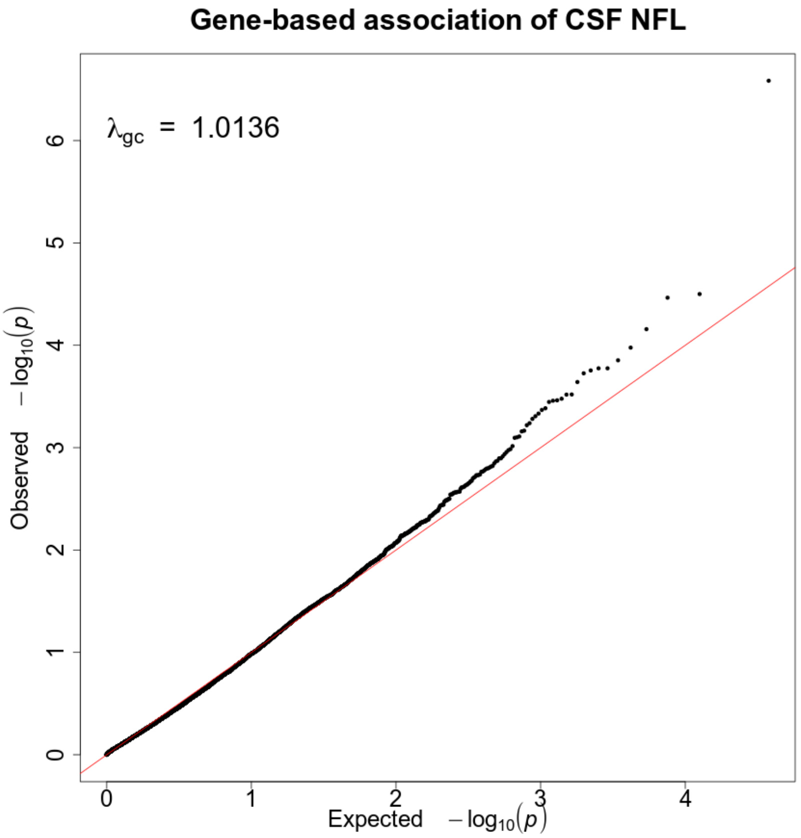

**Figure S2b.** log-transformed CSF_NF-L

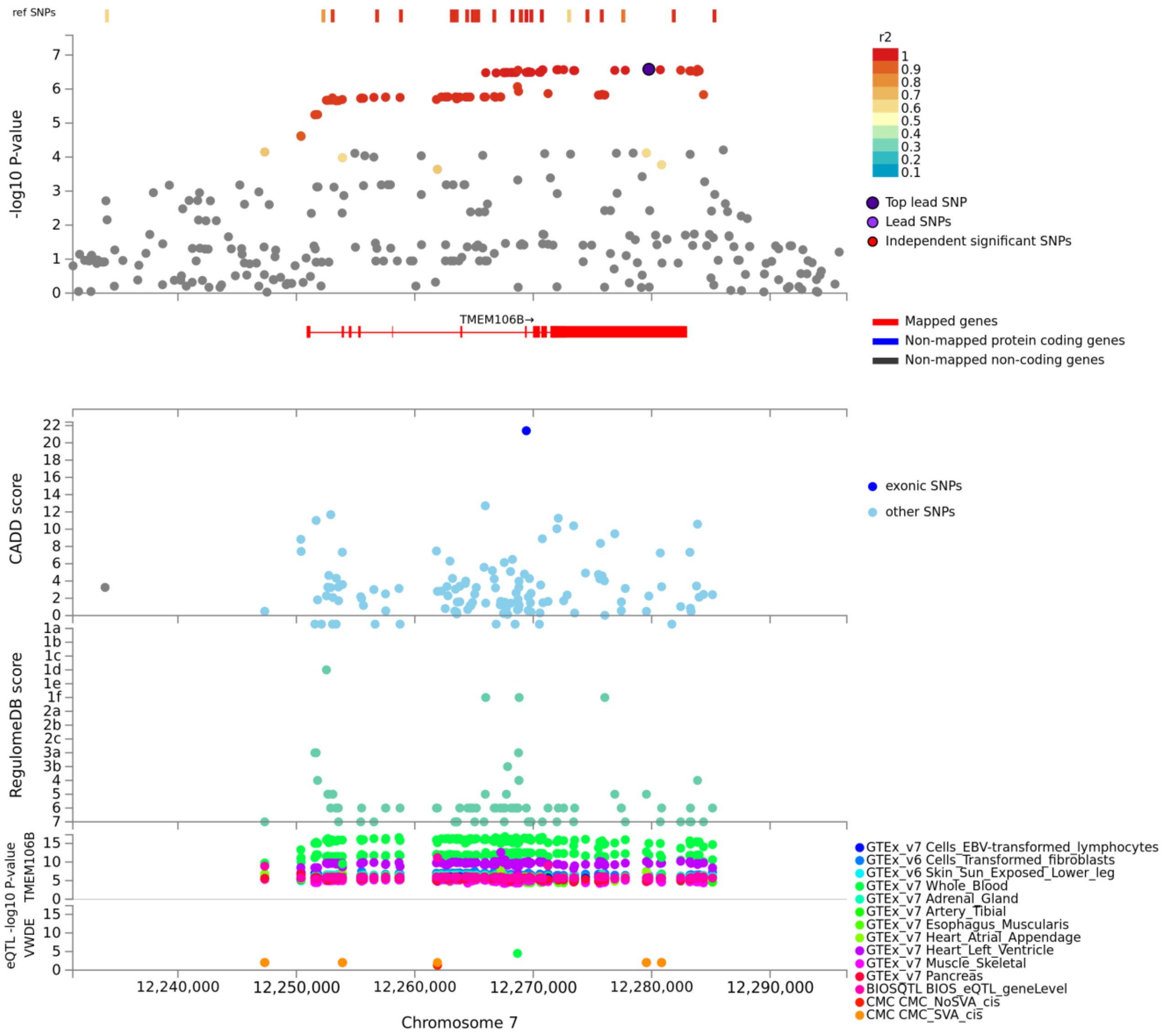

**Figure S3a.** log-transformed CSF_YKL-40

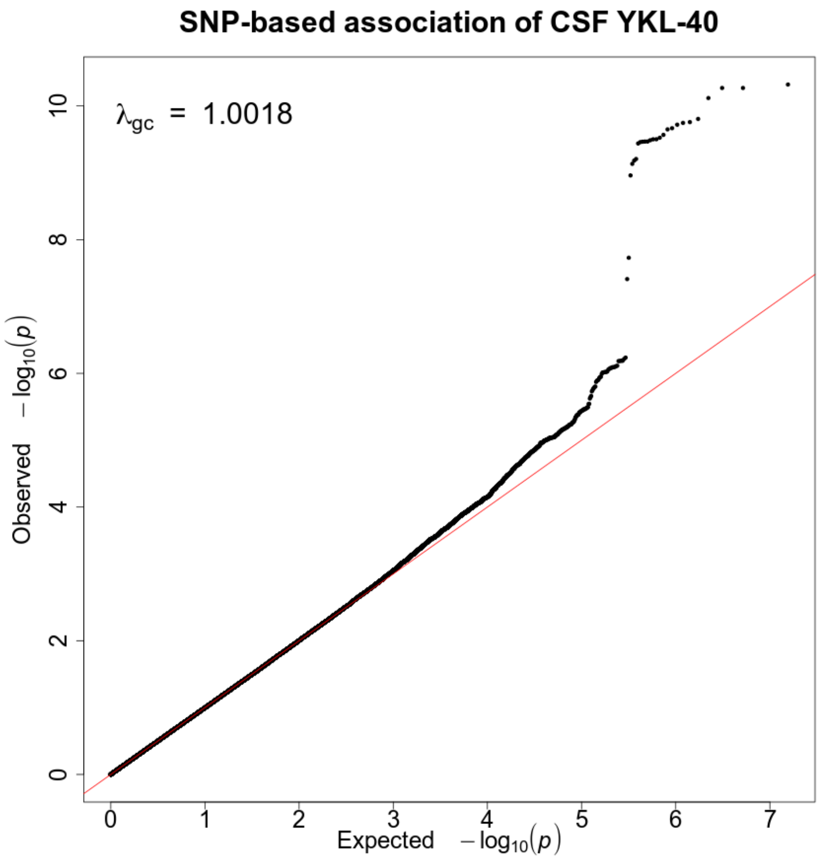


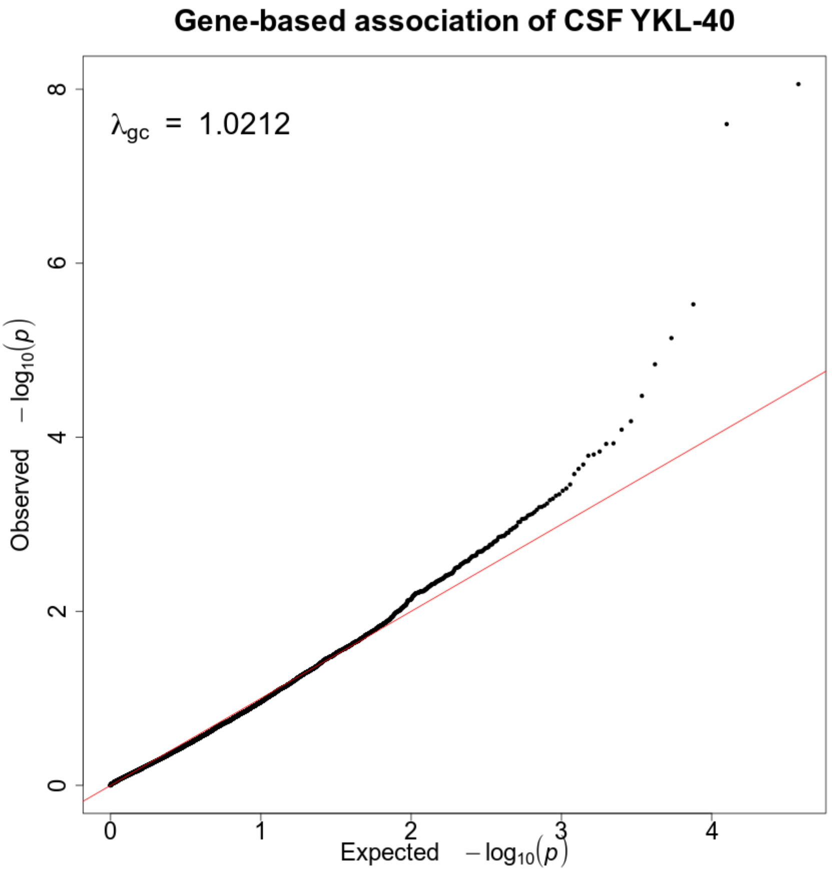

**Figure S3b.** log-transformed CSF_YKL-40

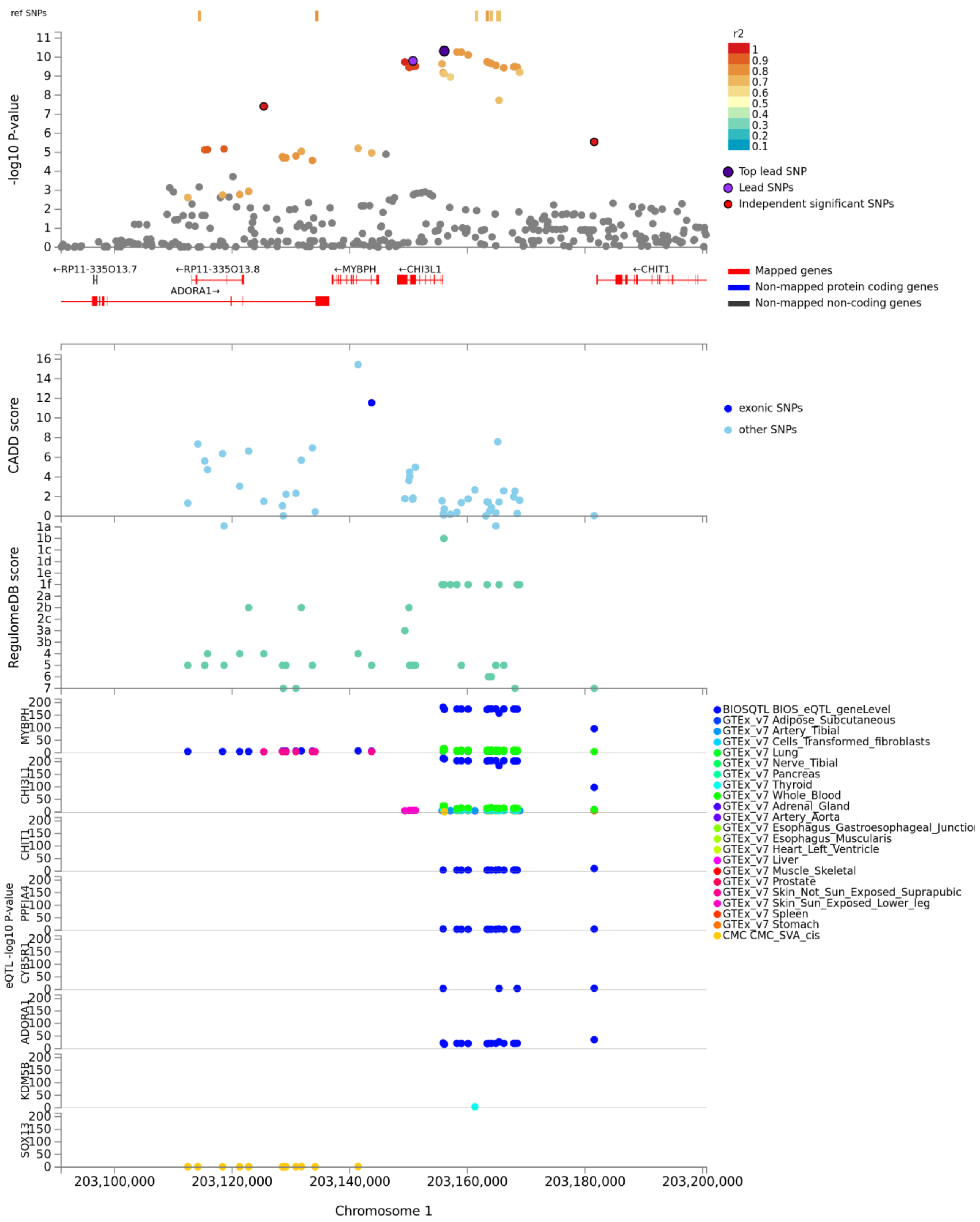

**Figure S4a.** log-transformed CSF_Ng

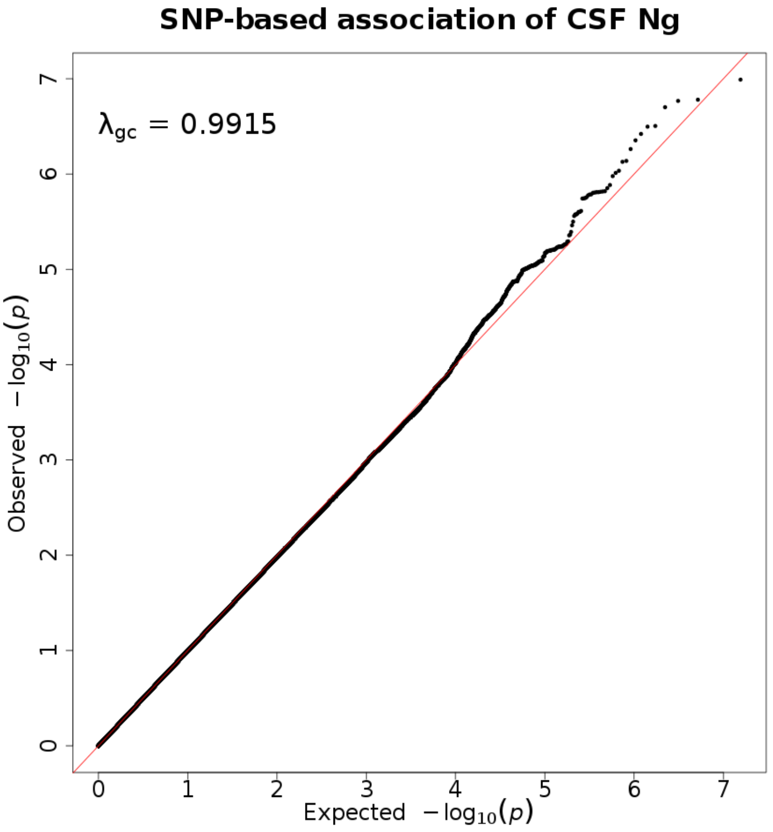


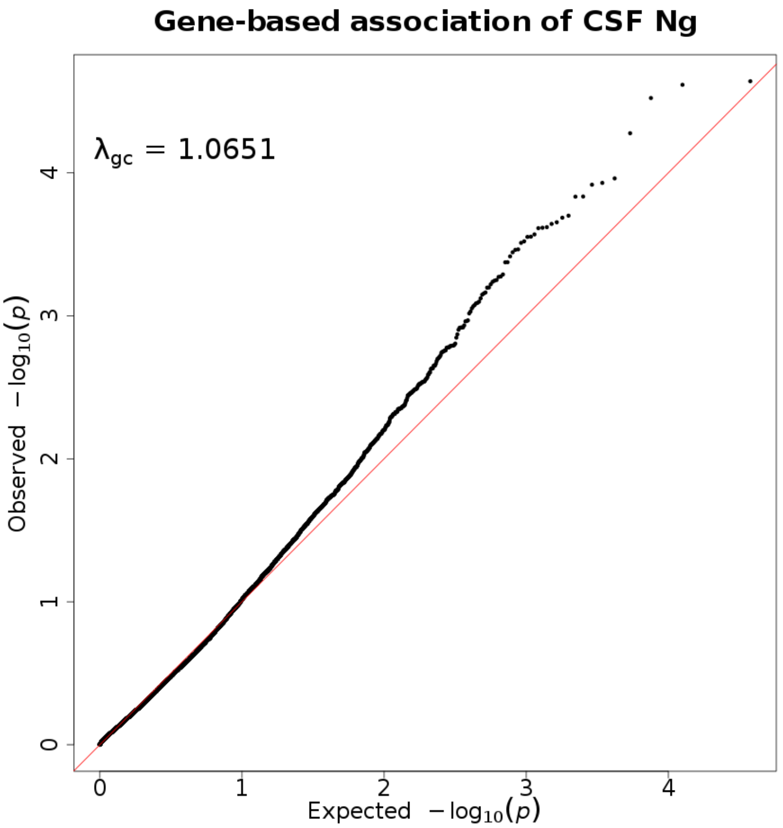

**Figure S4b.** log-transformed CSF_Ng

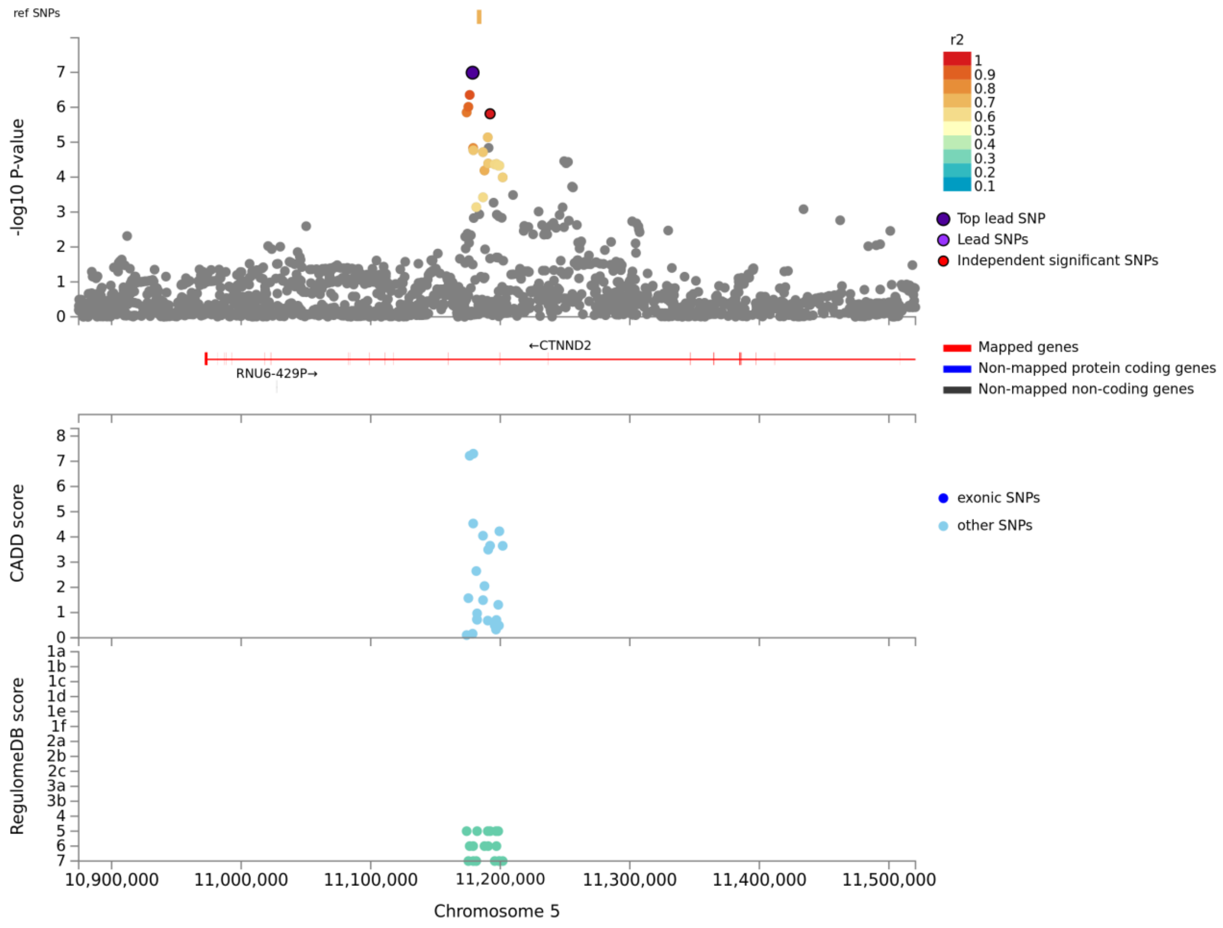
